# APOBEC2 is a Transcriptional Repressor required for proper Myoblast Differentiation

**DOI:** 10.1101/2020.07.29.223594

**Authors:** Jose Paulo Lorenzo, Linda Molla, Ignacio L. Ibarra, Sandra Ruf, Jana Ridani, Poorani Ganesh Subramani, Jonathan Boulais, Dewi Harjanto, Alin Vonica, Javier M. Di Noia, Christoph Dieterich, Judith B. Zaugg, F. Nina Papavasiliou

## Abstract

The activation induced cytidine deaminase/apolipoprotein B editing complex (AID/APOBEC) family comprises several nucleic acid editors with roles ranging from antibody diversification to mRNA editing. APOBEC2, an evolutionarily conserved member of this family, has neither an established substrate nor a mechanism of action, however genetic evidence suggests functional relevance in tissues such as muscle. Here, we demonstrate that in muscle, APOBEC2 does not have any of the attributed molecular functions of the AID/APOBEC family, such as RNA editing, DNA demethylation, or DNA mutation. Instead, we show that APOBEC2 occupies chromatin at promoter regions of certain genes, whose expression is repressed during muscle cell differentiation. We further demonstrate that APOBEC2 on one hand binds promoter region DNA directly and in a sequence specific fashion, while on the other it interacts with HDAC transcriptional corepressor complexes. Therefore, APOBEC2, by actively repressing the expression of non-myogenesis pathway genes, plays a key role in enforcing the proper establishment of muscle cell fate.

---

The AID/APOBEC proteins are zinc-dependent deaminases that catalyze the removal of the amino group from a cytidine base in the context of a polynucleotide chain, resulting in cytidine (C) to uridine (U) transition on DNA or RNA. Members of the AID/APOBEC family are closely related to one another based on homology and conservation of the cytidine deaminase domain containing a zinc-dependent deaminase sequence motif^1^. However, they differ by tissue-specific expression, substrates, and biological functions (reviewed in^2^). Physiologically these proteins alter the informational content encoded in the genome through a range of processes: acting as messenger RNA (mRNA) editors, affecting translation (APOBEC1)^3, 4^ acting as DNA mutators to create novel gene variants, restrict viruses and retrotransposons (AID and APOBEC3) (reviewed in^5^) and, changing DNA 5mC modification levels, leading to modulation of transcript abundance (AID and APOBEC1)^6, 7^.

APOBEC2 is an evolutionarily conserved member of the AID/APOBEC family^8^. Substantial evidence highlights the biological relevance of APOBEC2 in metazoans. In mice, APOBEC2 is highly expressed in cardiac and skeletal muscle where it affects muscle development^9^. Specifically, in the absence of APOBEC2, there is a shift from fast to slow muscle fiber formation, a reduction in muscle mass, and a mild myopathy with age^10^. In zebrafish, APOBEC2 has been implicated in muscle fiber arrangement^11^ and in retina and optic axon regeneration^12^. In frogs, APOBEC2 is important in left-right axis specification during early embryogenesis^13^. Mutations and gene expression changes of APOBEC2 have also been linked to cancer development^14, 15^.

Even though there is evidence for a biological role of APOBEC2, there are few insights to the mechanism by which APOBEC2 accomplishes these. Moreover, there has been no definite demonstration of its activity as a cytidine deaminase. Based on its homology with the other AID/APOBEC family members, it has been hypothesized that APOBEC2 may be involved in RNA editing^9, 14^ or DNA demethylation^6, 7, 12^. It has also been hypothesized that it has lost its deaminase activity altogether and may act biologically by a different mechanism^16^. However, there is currently a lack of knowledge on the direct physiological targets of APOBEC2, and its mechanism of action.

To answer some of these questions, we performed knockdown studies of APOBEC2 during the differentiation of the C2C12 murine myoblast cell line to systematically characterize the transcriptome and DNA methylation patterns of APOBEC2 deficient C2C12 cells. Our results confirm the requirement of APOBEC2 for myoblast to myotube differentiation. Additionally, we demonstrate the requirement of its amino-terminal disordered region for nuclear localization and myotube differentiation. While our results do not support APOBEC2 roles on RNA editing and on DNA methylation, we find that APOBEC2 downregulation leads to substantial gene expression changes affecting programs associated with myogenesis. Moreover, genomic occupancy experiments demonstrate that APOBEC2 interacts with chromatin at promoters of genes that are repressed during myoblast differentiation. Furthermore, these targets are derepressed with reduced abundance of APOBEC2, which allude to APOBEC2 acting as a transcriptional repressor. Notably, these target derepressed genes are not directly involved in myogenesis or muscle differentiation; instead they seem enriched for genes in the innate immune / inflammatory pathway. Finally, we show that APOBEC2 directly interacts with DNA as well as Histone Deacetylase 1 (HDAC1) repressor complexes, which supports the molecular function of APOBEC2 as a transcriptional repressor. Taken together, our data suggest that APOBEC2 has a direct role in regulating active gene transcription during myoblast differentiation as a transcriptional repressor.

## APOBEC2 is required for myoblast to myotube differentiation

The C2C12 myoblast cell line was derived from mouse satellite cells activated to proliferate after muscle injury in adult mice^17^. C2C12 myoblasts are thought to recapitulate the first steps of muscle differentiation in culture and upon differentiation induce APOBEC2 expression^13^ (Supplementary Fig. 1A), making them a suitable model to investigate putative roles of this suspected cytidine deaminase in situ.

To explore the role of APOBEC2 during myogenesis, we reduced APOBEC2 protein levels with short hairpin RNA (shRNA) against APOBEC2 mRNA. Protein reduction was efficient and coincided with reduced myoblast-to-myotube differentiation, evidenced by the decrease in expression of TroponinT and myosin heavy chain (MyHC), protein markers of late differentiation (Fig. 1A). At the cellular level downregulation of APOBEC2 protein levels coincided with reduced myotube formation (Fig. 1B). These observations match those previously reported using mouse embryonic stem cell-derived myogenic precursors^18^.

**Figure 1.**
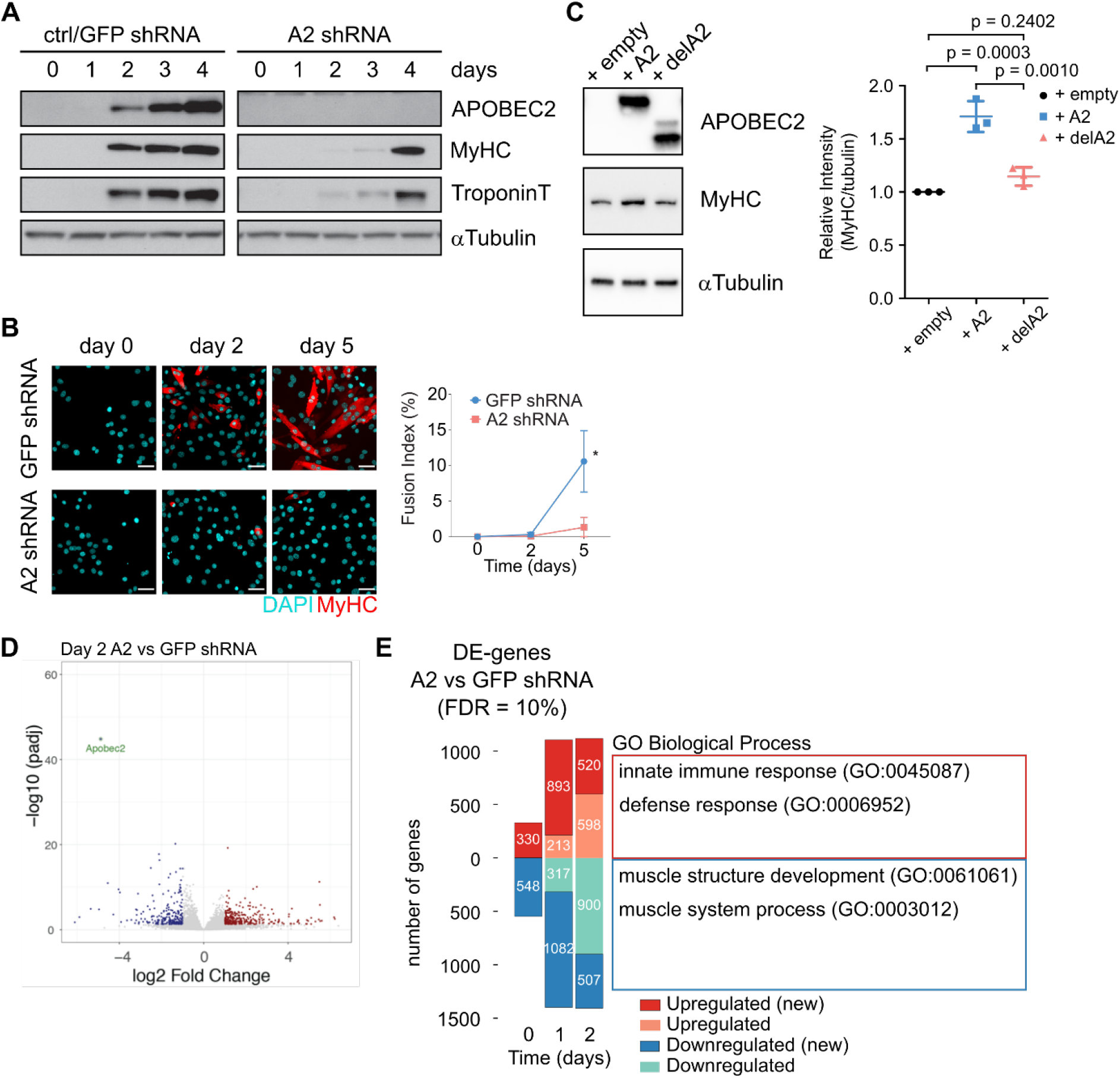
APOBEC2 expression is required for myoblast differentiation. **(A)** Cell lysates from C2C12 cell lines, ctrl/GFP shRNA and A2 shRNA, at different timepoints of myoblast differentiation (day 0 to day 4) were analyzed by Western blot. C2C12 myoblasts were transduced either with shRNA against GFP (ctrl/GFP shRNA) or shRNA against APOBEC2 (A2 shRNA). MyHC (myosin heavy chain) and TroponinT were used as markers of late differentiation; αTubulin, as loading control. **(B)** C2C12 cell lines were fixed and immunostained using an antibody specific to MyHC (red), DAPI (cyan) was used to stain for DNA. Scale bar = 50 μm. Line plot shows quantification of fusion index. Statistics: t test. Six fields of view were measured, and data is shown as means (n = 3). Error bars indicate SD; * = p < 0.05. **(C)** C2C12 knockdown cell line (A2 shRNA) was transduced with retrovirus overexpressing APOBEC2 or empty vector control (+ empty: transduced with empty vector, + A2: with APOBEC2 vector, and + delA2 with truncated APOBEC2). Cells were collected 96 hours post-transduction. Cell lysates were prepared and analyzed by Western blot. Representative blot from 3 independent biological replicates. Statistics: Ratio of MyHC mean intensity to αTubulin was normalized to corresponding + empty sample for each trial. Dot plot represent mean with error bars representing standard deviation (n = 3). One-way ANOVA with Tukey’s multiple comparisons test was performed to calculate p (adjusted P value). **(D)** Volcano plot was generated using log2 fold changes and p-adjusted values from comparing gene expression differences due to APOBEC2 knockdown at day 2 following induction of differentiation. Significantly differentially expressed genes with p adjusted value <0.05 are shown in red (upregulated) or blue (downregulated) and not significantly differentially expressed genes in gray. APOBEC2 is shown in green. **(E)** Number of differentially expressed genes (DE-genes) between APOBEC2 knockdown (A2 shRNA) relative to GFP shRNA control. Colors indicate up- (red) and down-regulated (blue) genes. A false discovery rate (FDR) cutoff of 10% was used to determine DE-genes. Dark red and blue indicate newly differentially expressed genes at a given time point. Common GO Biological Process terms across day 0 to 2 from statistical enrichment test ranking genes by log2FoldChange (see Supplementary File 1 for complete output tables of the tests).

We then restored APOBEC2 protein in these knockdown cells through overexpression. This led to an increase in MyHC levels, which confirms the essential role of APOBEC2 in myoblast differentiation in this model (Fig. 1C). Additionally, we produced truncated APOBEC2 (residues 41-224 mouse APOBEC2) based on its published structures^19, 20^. Truncated APOBEC2 (del(2-40)A2), was unable to restore MyHC expression levels. Interestingly, the truncated form of APOBEC2 was only found in the cytoplasmic fraction of differentiated C2C12 myoblasts (Supplementary Fig. 1C). We speculated that the nuclear localization of APOBEC2 was necessary for its role in muscle differentiation and pointed to a molecular function within the nuclear compartment.

## APOBEC2 loss leads to gene expression changes related to muscle differentiation

To study how APOBEC2 loss leads to problems in C2C12 myoblast-to-myotube differentiation, we performed RNA sequencing (RNA-Seq) to compare the transcriptome dynamics of APOBEC2 knockdown and control cells during differentiation. We observed that reduced APOBEC2 levels led to substantial gene expression changes (Fig. 1D,E). Notably, genes downregulated during differentiation were enriched for muscle development Gene Ontology (GO) terms, whereas genes upregulated were enriched for GO terms related to immune response (Fig. 1C; Supplementary File 1). The decreased expression of muscle differentiation related genes reflects the observed reduced myotube formation. Though undetectable on the immunoblot (Fig. 1A), perturbation of APOBEC2 levels prior to inducing differentiation seems to affect the potential of C2C12 to differentiate into myotubes. Genes involved in muscle development or function were also downregulated at day 0 prior to inducing the cells to differentiate.

Furthermore, genes related to cell development or differentiation GO terms, particularly immune system development, blood vessel development, and nervous system development, are among those overrepresented in the list of upregulated genes with APOBEC2 deficiency (Supplementary File 2). These spurious non-muscle transcriptional programs possibly reflect transdifferentiation events, which have been observed for C2C12 myoblasts^21^.

We next investigated possible molecular mechanisms of how APOBEC2 causes these gene expression changes. Due to the conserved cytidine deaminase domain within the AID/APOBEC family, APOBEC2 is posited to be an RNA editor^9, 14^ and a DNA demethylase^6, 7, 12^. Upon comparing the transcriptomes of the APOBEC2 knockdown and control C2C12 cells for instances of C-to-U RNA editing, using a previously validated pipeline^22^, we could not identify C-to-U or A-to-I RNA editing events that were APOBEC2 dependent (Supplementary Fig. 2A). Similarly, using bisulfite sequencing, we were unable to observe significant methylation differences between the APOBEC2 knockdown and control C2C12 cells that could account for the gene expression changes (Supplementary Fig. S2B-C). Altogether, these results strongly indicate that APOBEC2 is neither involved in mRNA deamination nor DNA demethylation in differentiating muscle.

**Figure 2.**
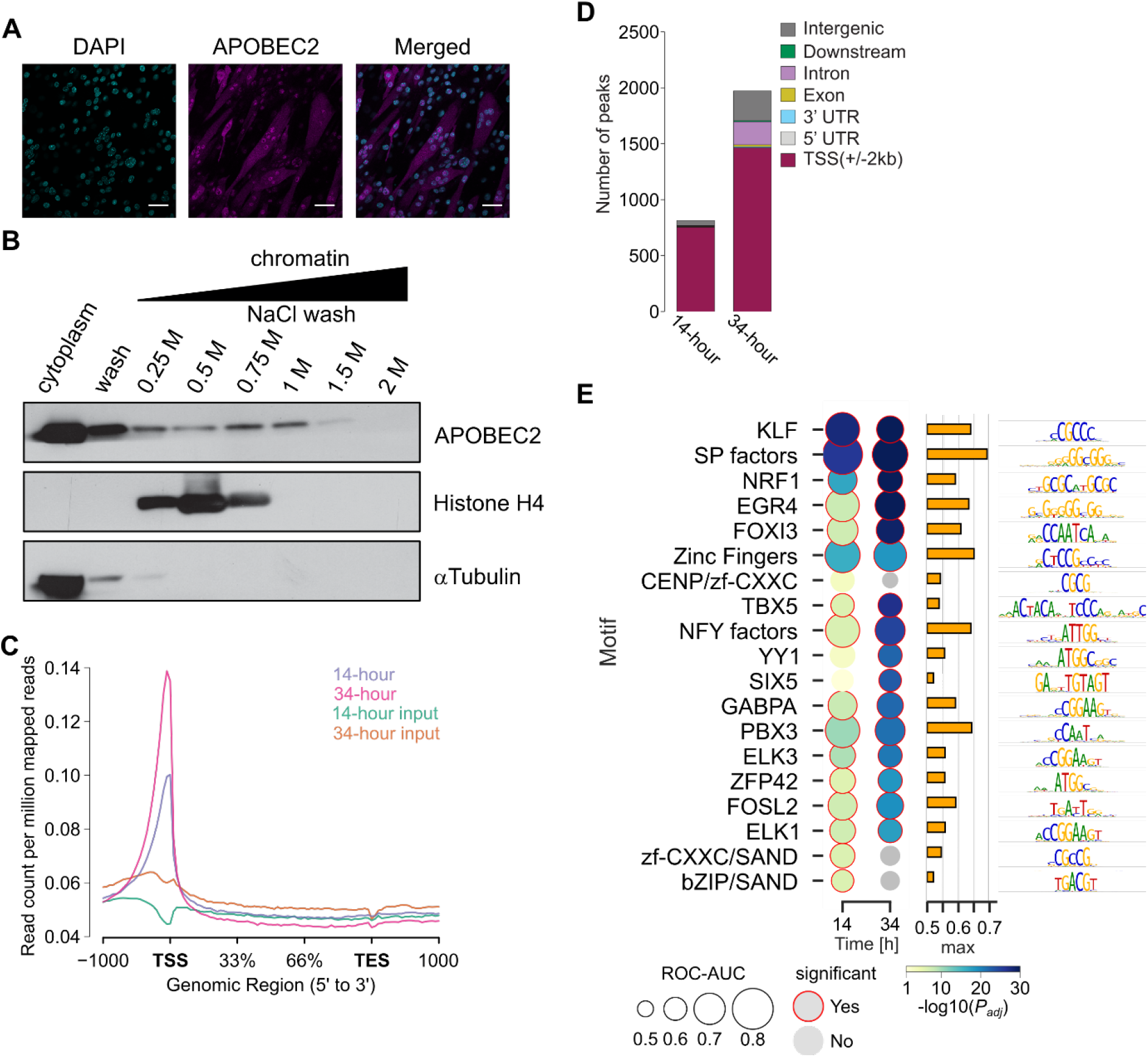
APOBEC2 localization and ChIP-Seq in differentiating C2C12 myoblasts. **(A)** Immunostaining of APOBEC2 (magenta) and DAPI-positive (blue) nuclei in differentiated C2C12 myoblasts (5 days in differentiation medium). Scale bar = 50 μm. **(B)** The sequential salt extraction profile of endogenous APOBEC2 and histone H4 are shown. alpha-tubulin is a cytoplasmic marker. The amount of indicated proteins in eluates was measured by Western blotting. **(C)** The mean normalized APOBEC2 signal (plotted as read counts per million mapped reads) across all annotated genes. This plot shows the global differences in APOBEC2 binding between the two time points in TSS. Both time points are in biological triplicates. **(D)** Genomic annotations of APOBEC2 consensus binding regions in each of the time points (14- and 34-hour). Binding regions are annotated based on genomic feature. The priority of assignment to a genomic feature when there is annotation overlap is: Promoter (2kb around the TSS), 5’ UTR, 3’ UTR, Exon, Intron, Downstream (within 3kb downstream of the end of the gene). **(E)** Enrichment of 8-mers associated to 19 known transcription factors specificities (motifs) in APOBEC2 ChIP-seq regions, compared against negative genomic regions (see Methods). Receiver Operating Characteristic Area under the Curve (ROC-AUC, circle sizes) values are descendently sorted on the y-axis by minimum adjusted P value observed in both time points (color). All 18 shown transcription factors are significant on at least one time point (adjusted P value < 0.1, using one-sided Wilcoxon rank sum test and Benjamini-Hochberg correction. Red circles). Right barplot indicates maximum ROC-AUC value, row-wise. Right sequence logos depict unsupervised alignment of all 8-mers referred to that transcription factor motif^29^.

## APOBEC2 binds promoter regions

Cytidine deaminases of the AID/APOBEC family can bind and mutate DNA either at gene bodies, e.g. exons of the immunoglobulin locus, as catalyzed by AID, reviewed in^23^ or at promoter regions, e.g. local hypermutations as catalyzed by APOBEC3 family members - reviewed in^24^. To assess whether APOBEC2 could also bind genomic DNA, we first determined the subcellular localization of APOBEC2 in muscle cells. We observe that APOBEC2 is present in both the cytoplasm and nucleus of differentiated C2C12 myotubes (Fig. 2A, Supplementary Fig. 1C). A weak nuclear localization signal (NLS) can be predicted by cNLS Mapper^25^ (APOBEC2 residues 26 to 57, with a score of 3.7) that could explain the lack of nuclear localization of truncated APOBEC2, del(2-40)A2. However, full length APOBEC2 does not show NLS activity but is homogenously distributed throughout the cell, presumably through passive diffusion^26^. To then assess whether nuclear APOBEC2 could bind chromatin, we utilized sequential salt extraction based on the principle that loosely bound proteins will be dissociated at low salt concentration, while tightly bound ones will not^27^. Using this technique, we found that a fraction of APOBEC2 within differentiating C2C12 cells, remains bound to chromatin even at high salt concentrations (up to 1 M NaCl). As a comparison histone H4 dissociates completely from DNA at 0.75 M NaCl (Fig. 2B). These data suggest a strong association of nuclear APOBEC2 with chromatin in differentiating myoblasts.

To determine APOBEC2 binding localization within chromatin, we performed chromatin immunoprecipitation-sequencing (ChIP-Seq) experiments, and calculated enrichment of APOBEC2 at specific loci over input using MACS2 (see Methods). We performed each experiment in triplicate, and only peaks that were called in at least 2 out of 3 replicates were analyzed. Importantly, we queried APOBEC2 occupancy at two different time points, 14- and 34-hours post-differentiation, that precede the RNA-Seq time points, where we observed changes in gene expression and represent time points of low and higher APOBEC2 protein abundance. Overall, the signal around peak summits of transcription start sites (TSS) is higher at 34 versus 14 hours, reflecting an increase of APOBEC2 in chromatin. In contrast, input controls show lower enrichment over the peak summits (Fig. 2C).

Annotating the APOBEC2 peaks by genomic feature showed that for both time points most of the peaks fall within promoters, defined as regions −2 kilobases (kb) to +2 kb around the TSS (Fig. 2D). Next, we wanted to determine whether APOBEC2 occupies specific motifs within promoter regions. The members of the AID/APOBEC family that bind to DNA have some local sequence preference with regard to sites they mutate, but do not display rigorous sequence specificity, e.g. akin to transcription factor (TF) binding sites^28^. To assess the motif signatures in those regions we queried through motif enrichment the overrepresentation of TF 8-mers sequences associated to main TF modules^29^. Among 108 TF modules tested against a controlled background of negative sequences (see Methods), we observed 19 of those significantly enriched at APOBEC2 regions in at least one differentiation time point. SP and KLF motifs were among the top enrichments observed (Fig. 2E), suggesting a co-regulatory role between these factors and APOBEC2. In general, TF specificity groups increase their significance between 14 and 34 hours but with lower overall effect sizes or only at the later time points, suggesting that the interplay between TFs and APOBEC2 occupied regions is specific at 14 hours but broader at later time points, likely as consequence of events related to its early binding. Additionally, we did not observe APOBEC2 related DNA mutation at the occupied peaks, indicating that APOBEC2 is not a DNA mutator like other AID/APOBEC family members (Supplementary Fig. 3).

## APOBEC2 represses expression of occupied genes

Focusing on the promoter bound genes we determined that there are ~1500 genes that are bound by APOBEC2 near their transcription start sites in any of the time points, 590 of which are occupied at both time points (Fig. 3A). Overall, about 79% of the genes that are bound by APOBEC2 in the 14-hour time point remain bound at the 34-hour time point. Further, using Gene Set Enrichment Analysis to determine the distribution of APOBEC2 occupied genes at both time points in a list of ranked expression changes either through differentiation (day 0 to 2) or upon APOBEC2 knockdown (A2 shRNA vs GFP shRNA at day 2). Our results show that APOBEC2 occupied genes are significantly enriched at genes downregulated through differentiation (Fig. 3B). Moreover, upon APOBEC2 knockdown, APOBEC2 occupied genes are instead enriched at upregulated genes (Fig. 3C). Interestingly, overall expression of APOBEC2 bound genes through C2C12 differentiation from day 0 to 2 is significantly decreased during differentiation compared to genome wide expression changes (−0.3 versus 0.1 mean log2 fold changes; *t* = −8.2; adjusted P-values < 0.0001; Fig. 3D). Furthermore, expression of APOBEC2 occupied genes increase upon APOBEC2 knockdown (*t*-statistic = 4.0; adjusted P-value < 0.001) highlighting a global repressive role of APOBEC2 during differentiation. Altogether, these results suggest that the observed transcriptional changes linked to APOBEC2 deficiency are due to APOBEC2 functioning as a transcriptional repressor.

**Figure 3.**
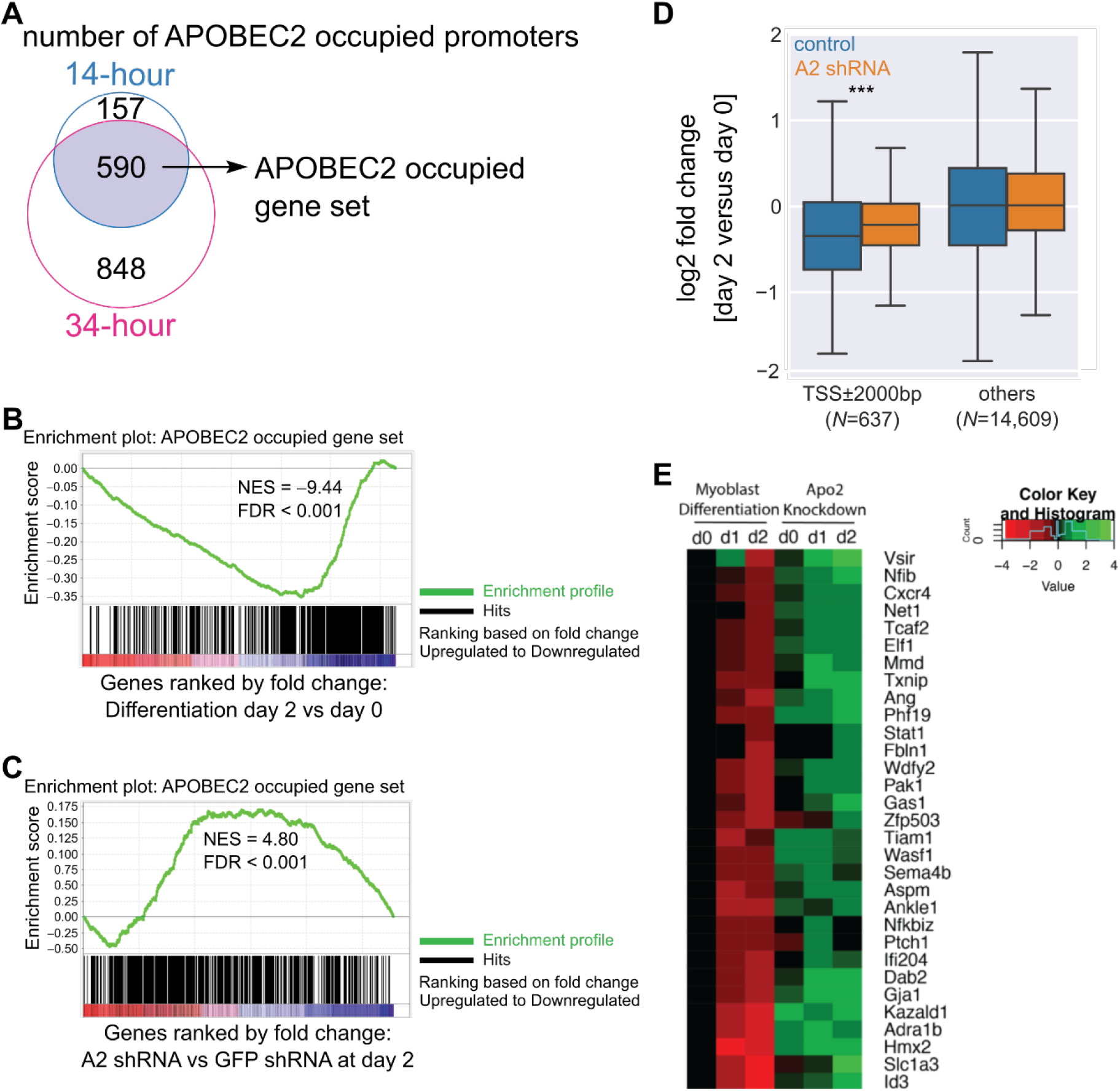
APOBEC2 binding at specific transcription factor DNA motifs. **(A)** The Venn diagram represents the number of genes that have APOBEC2 occupancy on their promoters at 14- and 34-hour time points. The genes that show consistent APOBEC2 occupancy in their promoters at both time points were used to create an APOBEC2 occupied gene set as shown in the Venn diagram. **(B,C)** Gene set enrichment analysis (GSEA) (Subramanian et al., 2005) was used to test the enrichment of the APOBEC2 occupied gene set in the list of genes that are differentially expressed through differentiation (C) or the ones that are differentially expressed due to APOBEC2 knockdown at day 2 (D). The enrichment profile over the whole ranked gene set is shown in green with normalized enrichment score (NES) and FDR values. Gene hits are shown as black lines. A positive NES indicates gene set enrichment at the top (positive/up fold change) of the ranked list; a negative NES indicates gene set enrichment at the bottom (negative/down fold change) of the ranked list. **(D)** Expression changes for genes in control (GFP control shRNA) and A2 knockdown (A2 shRNA) C2C12 during differentiation. Genes are grouped by the presence of an APOBEC2 ChIP-seq peak nearby Transcription Start Sites (TSS) at not more than 2000bp from it, or genome wide background (others). Asterisks indicate adjusted P values, by two-sided t-test corrected using Benjamini Hochberg procedure (***=0.001). **(E)** Heatmap showing expression changes of genes related to differentiation (regulation of differentiation, GO: 0045595) occupied by A2 during differentiation in C2C12 cells and A2 knockdown cells. The list of APOBEC2 repressed genes were filtered from the C2C12 RNASeq data of A2 occupied genes (defined as genes with A2-ChiPSeq peak in the promoter region). Genes repressed during differentiation log2 fold change greater than 0.58 (absolute fold change > 1.5) at day 1 or 2 of differentiation and upregulated in the knockdown (versus control) at day 0, 1, or 2 were selected. These genes were then used as input for statistical overrepresentation test – GO biological processes through Panther (ver 14) with the default settings using all *M. musculus* genes in database as reference list^81^.

To validate its possible role as a transcriptional repressor, we selected candidate genes repressed by APOBEC2 occupancy. We narrowed it down to a list of genes occupied by APOBEC2 which are downregulated during differentiation (day 2) and upregulated with APOBEC2 knockdown. In this gene list, we did not find an overrepresentation of GO terms relating to muscle differentiation; but rather, we found terms related to development of other lineages (Supplementary Table S1) similar to the non-muscle genes upregulated with APOBEC2 deficiency. Furthermore, a list of 31 genes related to differentiation (GO: 0045595 regulation of differentiation) in other tissue contexts were upregulated with APOBEC2 loss (Fig. 3E). We speculate that APOBEC2 is acting as a repressor to direct C2C12 differentiation into the muscle-lineage by repressing specious gene networks related to other lineages.

## APOBEC2 binds directly to single- and double-stranded DNA

Thus far, the results suggest that APOBEC2 has a molecular role unique to the AID/APOBEC family. While it does not have the capacity to modify nucleic acids (RNA or DNA) through deamination^30, 31^, it is capable of binding chromatin to regulate transcription. This implies either that APOBEC2 interacts directly with DNA at promoter regions, or that it interacts with transcription regulators that do so, or both.

To assess whether APOBEC2 is capable of binding DNA directly, we generated recombinant APOBEC2 (expressed in insect cells, Supplementary Fig. 4A) and tested it in vitro for its capacity to bind DNA oligonucleotides that represent APOBEC2 bound sequences in vivo. We tested an SP/KLF DNA motif as it was highly represented in the ChIP-Seq data (Figure 4A). As a negative control, we tested an A-tract motif that based on the ChIP-Seq was not bound by APOBEC2 but which occurred frequently at promoters of differentially regulated genes. Using microscale thermophoresis (MST), purified APOBEC2 bound both single-stranded and double-stranded DNA containing the SP/KLF motif with reasonable affinity (calculated Kd = 253 ± 86 nM and 709 ± 186 nM, respectively, Figure 4A, S4B). In contrast, measurements for the A-tract motif showed large variances between trials, indicating lack of specific binding (Figure S4C). This ability to selectively bind both ssDNA and dsDNA in a sequence-specific manner in vitro (Fig. 4A) and in vivo (Fig. 2E) makes APOBEC2 unique among AID/APOBEC family members. These results demonstrate that APOBEC2 can target specific genes at promoter regions, and through this binding repress transcription.

**Figure 4.**
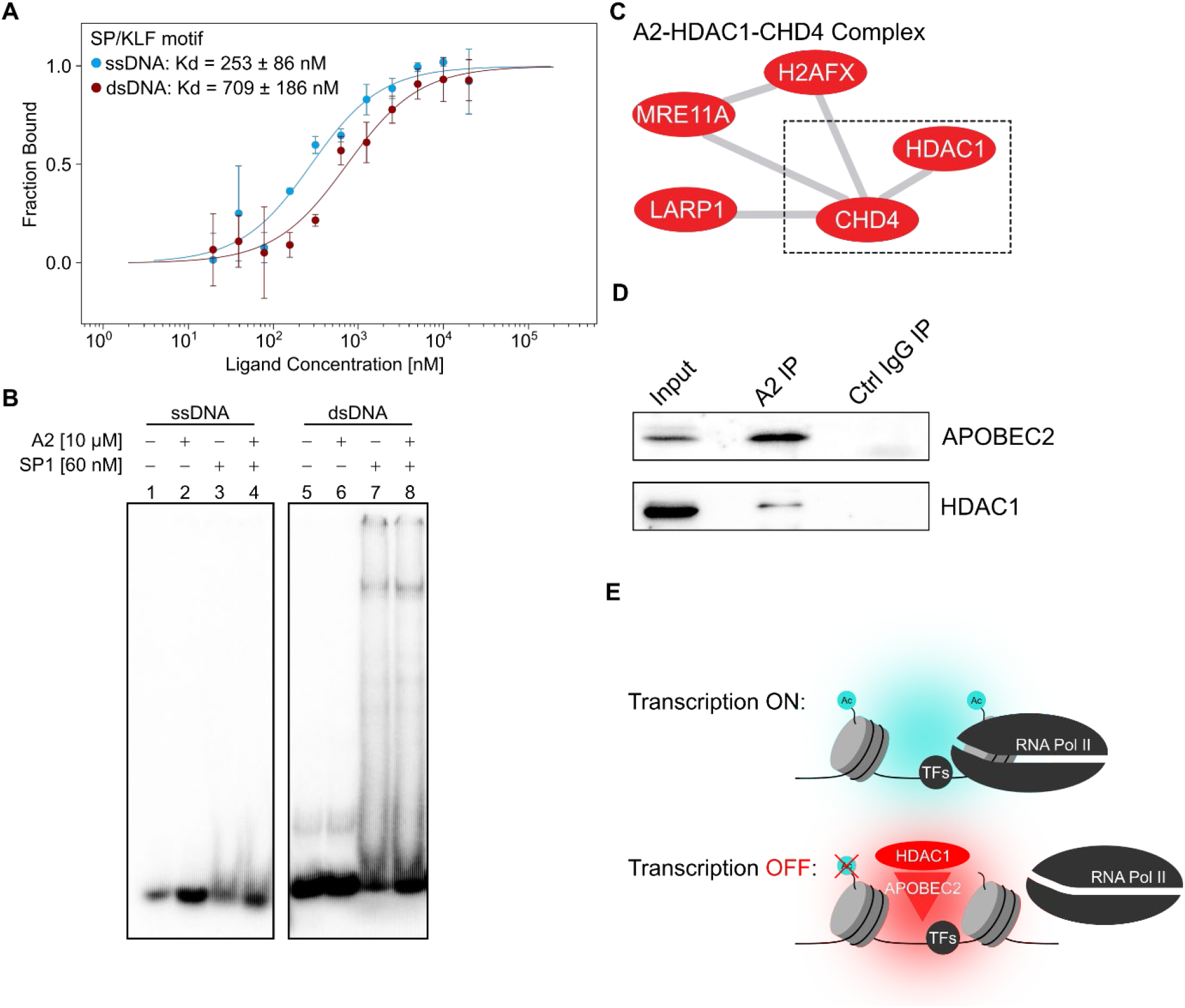
APOBEC2 recruits the HDAC1 co-transcriptional repressor complex. **(A)** Microscale thermophoresis (MST) experiments measuring purified APOBEC2 binding to single-stranded DNA (ssDNA) or double-stranded (dsDNA) SP/KLF motif. Cy5-labeled APOBEC2 was kept constant (50 nM) while the concentration of non-labeled SP/KLF motif was titrated (1:1 dilution) between 20 nM – 20,000 nM. The calculated Kd was computed using the standard settings (thermophoresis + T jump) with the MO.Affinity Analysis v2.1 (NanoTemper Technonolgies). Values represent 3 independent measurements with error bars representing the standard deviation. **(B)** Electrophoretic mobility shift assays (EMSA) of recombinant mouse APOBEC2 and human SP1 protein on either ss or dsDNA with an SP/KLF motif. Radioactively labeled ss or dsDNA (1 nM) was mixed with either recombinant APOBEC2 (10 uM) or SP1 (60 nM) protein, or both. Gel shift image is representative of at least 3 independent experiments. **(C)** Selected protein complex identified by APOBEC2 proximity-dependent protein biotinylation (BioID). Each red node corresponds to a protein that was identified by BioID mass spectrometry to interact with APOBEC2. CHD4 was also identified in a BioID data comparing APOBEC2 and AID in mouse B cells^82^. The edges denote the known interactions of these proteins with each other (see Figure S5B for other complexes). **(D)** Co-immunoprecipitation (Co-IP) of APOBEC2 with HDAC1 in C2C12 myoblasts differentiated to myotubes for 4 days. Nuclear protein lysates (Input) were incubated with beads conjugated to either APOBEC2 antibody (A2 IP) or IgG isotype control antibody (Ctrl IgG IP). Proteins were then eluted, ran on an SDS-PAGE gel, and blotted with APOBEC2, or HDAC1 antibodies. **(E)** Proposed molecular function of APOBEC2 as a co-transcriptional repressor complex that acts on active/open chromatin to repress transcription through HDAC1 histone deacetylation during myoblast differentiation.

Using gel electromobility shift assays (EMSA), we checked for APOBEC2 cooperativity with the transcription factor SP1 on its cognate motif. As expected, purified SP1 protein alone shifted dsDNA with the SP/KLF motif (lane 7, Fig. 4B). Unexpectedly, we were unable to see a shift by APOBEC2 alone on either ssDNA or dsDNA unlike with MST (lanes 3 and 6, Fig. 4B). This could be due to the running conditions which were unable to preserve the presumably weaker intermolecular affinity. However, APOBEC2 together with SP1 produced a shift with stronger intensity indicating cooperativity between the two DNA binding proteins (lane 7 vs lane 8, Fig. 4B). Interestingly, the two do not interact directly (Supplementary Fig. 4D), suggesting that the enhanced SP1 binding is mediated through DNA – possibly through changes in DNA conformation given that APOBEC2 has no detectable deaminase activity. This cooperativity suggests a molecular mechanism wherein APOBEC2 alters transcription factor affinity for specific motifs leading to transcriptional repression.

## APOBEC2 interacts with corepressor complexes in vivo

To ascertain the mechanism of action for the observed transcriptional repression, we used proximity-dependent protein biotinylation (BioID) to confirm and identify other putative APOBEC2 proximal protein complexes that may indicate direct interactions. After statistical curation we identified 124 proteins that were significantly more tagged by APOBEC2-BirA and/or BirA-APOBEC2 than GFP-BirA controls in C2C12 cells (Supplementary Table S2). Functional annotation showed that many APOBEC2 neighboring proteins are related to cell membrane and cytoskeleton organization processes, in line with its high cytoplasmic abundance (Fig. 2B), but terms related to chromatin modification and histone deacetylation were also enriched (Supplementary Fig. 5A and Supplementary Table S3). Of special interest was the identification the histone deacetylase 1 (HDAC1) and Chromodomain-helicase-DNA-binding protein 4 (CHD4), both components of the nucleosome remodeling and histone deacetylation (NuRD) transcriptional corepressor complex^32^, (Fig. 4C and Supplementary Fig. 5B). Using co-IP, we validated that APOBEC2 interacts with HDAC1 in differentiated C2C12 myoblasts (Fig. 4D). Together with the observation that APOBEC2 interacts directly with chromatin, this suggests that APOBEC2 plays a role in gene regulation through epigenetic nucleosome modification with these transcriptional corepressor complexes.

Taken together, our results suggest a direct role of APOBEC2 in repressing specific transcribed genes, likely mediated by an HDAC1 co-repressor complex during C2C12 myoblast differentiation (Fig. 4F).

## DISCUSSION

There have been many hypothesized roles for the cytidine deaminase APOBEC2. Here we show that the expression of APOBEC2 during myoblast differentiation has consequential effects on myotube formation owing to at least one unexpected molecular function: transcriptional control. We discovered that APOBEC2 loss, leads to faulty myoblast differentiation and concomitant gene expression changes. We show that these gene expression changes come about through direct chromatin interaction of APOBEC2 at promoters – with no observed APOBEC2 related changes in RNA editing, DNA methylation, or DNA mutation. We find instead that APOBEC2 is capable of directly binding DNA and interacting with corepressor complexes. Indeed, through this interaction, APOBEC2 specifically represses transcriptional programs unrelated to muscle differentiation thus indirectly supporting proper muscle differentiation.

The deaminase domain of APOBEC2 appears to have lost catalytic activity^30, 31^. However, it seems to have retained its ability to bind nucleic acids. Binding ssDNA is nothing new to members of the AID/APOBEC family^33^. However, the demonstration that APOBEC2 directly binds to dsDNA is groundbreaking for the family. There has been evidence for APOBEC3-driven dsDNA mutation but no evidence for direct dsDNA binding; instead, the mutational mechanism is likely through ssDNA^34, 35^. Prior work has linked APOBEC2 overexpression to RNA editing of specific transcripts as observed in the healthy livers of transgenic mice which eventually develop hepatocellular carcinoma^14^. Notably, RNA editing was only detectable in the liver at specific transcripts for the transgenic mice. However, based on our transcriptome analysis, we were unable to find evidence of such RNA editing in our myoblast differentiation models. Prior work has also reported mild effects of APOBEC2 on DNA methylation specifically at the MYOG promoter^18^; yet from our ChIP-Seq data, we do not find occupancy at the MYOG gene. Furthermore, there is conflicting data on the role of the AID/APOBEC family in active DNA demethylation; and, APOBEC2 dependent demethylation has not been found in other cellular contexts^36, 37^. A role in transcriptional repression such as we propose, would also explain these prior data, as indirect effects of improper differentiation.

Previous studies suggested that recombinant APOBEC2 is incapable of binding DNA or deaminating it^16, 30, 31^. Our experiments using recombinant protein produced in eukaryotic cells show that APOBEC2 is capable of binding DNA with reasonable affinity, and when it does so, it alters the ability of proximal transcription factors (such as SP1) to interact with their cognate motifs. Similar findings have also been observed with the transcription factor POU6F2, suggesting but not proving a role for APOBEC2 in transcription^16^.

Furthermore, our proximity ligation experiments reveal that APOBEC2 protein directly interacts with HDAC1 and CHD4 containing repressor complexes providing a direct epigenetic mechanism for the repression of APOBEC2 occupied genes. HDAC1 and CHD4 are components of the NuRD corepressor complex, whose involvement has already been proposed in skeletal or myocardial muscle fate determination^38^. Additionally, the abundance of APOBEC2 in the cytoplasm and its proximity to proteins that participate in cell morphology and the cytoskeleton could either reflect a mechanism limiting APOBEC2 access to the nucleus, or perhaps an additional function that remains to be studied.

Uniquely amongst AID/APOBEC family members, the amino-terminus of APOBEC2 contains a region that is glutamate-rich and intrinsically disordered^20^. Loss of this unstructured domain results in inability of the protein to rescue the knockdown phenotype likely through the loss of its nuclear retention. Proteins with similar disordered regions have been shown to form liquid phase separated membrane-less compartments and this function appears to be especially important for transcriptional regulation as transcription factors with disordered regions have been shown to form such compartments^39^. Even though, no enzymatic activity has been detected for APOBEC2, potential substrates could be present in these chromatin compartments: transient RNA species, eluding detection limits by our RNA-Seq method, or, potentially, ssDNA structures at melted promoters. We propose that through its N-terminal unstructured region APOBEC2 is retained in the nucleus where on one hand it binds DNA at promoter regions in a sequence specific fashion, and on the other, it recruits corepressor complexes to repress transcription.

We hypothesize that APOBEC2 acts as a modulator of its bound promoters during the myogenic program – fine-tuning it for muscle differentiation and repressing other lineage programs. MYOD1 has been shown to bind and activate lineage programs outside the muscle lineage; however, this is mitigated by corepressors^40^. Furthermore, as APOBEC2 expression under healthy conditions is mostly confined to muscle tissue (both skeletal and heart), where it might be acting as a ‘many-but-muscle’ lineage repressor – similar to MYT1L in neuronal differentiation^41^.

The discovery that APOBEC2 has a direct role in transcriptional control impacts how we interpret the phenotypes that have been attributed to it in the mouse knockout models and other biological systems, well beyond muscle differentiation – for example zebrafish retina and optic nerve regeneration, *Xenopus* embryo development, and cancer development^12–14^. In the zebrafish models, APOBEC2 loss leads to similar defective muscle phenotypes but it is deemed essential in the retinal regeneration model – where cellular reprogramming is a key step^12^. Directly or indirectly, these prior observations likely relate to aberrant transcriptional programs, normally silenced in the context of tissue development or cell differentiation due to APOBEC2 transcriptional control. Taken together, we postulate that APOBEC2 is a transcriptional repressor that modulates transcriptional programs during cell differentiation or reprogramming.

## Methods

### Data availability

High throughput sequencing datasets are all found in: GSE117732 and more specifically: RNA-Seq (GSE117730); ChIP-Seq (GSE117729); ERRBS (GSE117731). Mass spectrometry data for BioID performed in Flp-In 293 T-REx cells have been deposited in MassIVE under ID.

### C2C12 cell culture

C2C12 cells (CRL-1772, ATCC) were maintained in DMEM (30-2002, ATCC) with 10% fetal bovine serum and fed every two days. To differentiate equal number of cells (2.5 x 10^5) were seeded in 6-well plates followed by media change to DMEM with 2% horse serum after 12 hours. For generating single cell clones for RNA-Seq and RRBS experiments C2C12s were sorted using fluorescence-activated cell sorting (FACS) and seeded into a 96 well plate. Each clone was expanded and tested for successful knockdown through immunoblotting.

### APOBEC2 knockdown and overexpression

C2C12 cells were infected with lentiviruses carrying shRNA, targeting either APOBEC2 or GFP. All APOBEC2 shRNAs were obtained from The Broad Institute’s Mission TRC-1 mouse library and present in pLKO.1-puro construct. Plasmids used: pLKO.1 - TRC cloning vector (Addgene, # 10878)^42^; pLKO.1 puro GFP siRNA (Addgene, # 12273)^43^. The design of shRNAs and cloning in pLKO.1-TRC, were done according to the Addgene protocol (Protocol Version 1.0. December 2006). The following shRNAs sequences were used for APOBEC2 knockdown: A2 shRNA: GCTACCAGTCAACTTCTTCAA; GFP shRNA: GCAAGCTGACCCTGAAGTTCAT.

Virions were produced by co-transfection of pLKO.1-puro shRNA containing construct, packaging plasmid psPAX2 (Addgene, #12260) and envelope plasmid pMD2.g (Addgene, #12259) in 293T cells (CRL-3216, ATCC). Transfections were done using Lipofectamine 2000 Reagent (Invitrogen) as per manufacturer instructions. Supernatants with lentiviral particles were collected at 24 and 48 hours after transfection, passed through a 0.45 mm filter and applied to C2C12s. For APOBEC2 constitutive knockdown, C2C12 cells were infected with pLKO.1 containing lentiviruses in growth media containing 8 μg/mL polybrene for 12 hours. Two days after lentiviral infection cells were cultured with 4 μg/ml puromycin containing media for two more days to select cells stably expressing the shRNA.

Rescue constructs, mouse APOBEC2 and del(1-41)APOBEC2 with silent mutations to escape shRNA knockdown, and tagged constructs, APOBEC2 and del(1-41)APOBEC2 with C-terminal 3xHA-tags, were cloned into pMXs-GFP/Puro retroviral vectors (Cell Biolabs, Inc.). Virions were produced in 293T cells by co-transfection with pMXs construct and pCL-Eco (Novus Biologicals) using Lipofectamine 2000.

### Immunoblotting and co-immunoprecipitation

For immunoblotting experiments, C2C12 cells were first washed with cold PBS and lysed in 100 μl RIPA lysis buffer (Santa Cruz, sc-24948) in 6-well plates. They were incubated at 4°C for 15 minutes and then extracts were scrapped into a microfuge tube. Lysates were snap frozen in liquid nitrogen. After thawing the lysates on ice and clearing out cell debris by centrifugation, equal amounts of total protein (ranges between 10-30 μg) were boiled in SDS-PAGE sample buffer and loaded onto each lane of a polyacrylamide gel (Criterion XT Bis-Tris Gel 12%, Bio-Rad). Following electrophoresis, the resolved protein was transferred to a nitrocellulose membrane and subjected to western blot analysis. The source and dilution for each antibody used were: polyclonal rabbit-APOBEC2 (gift from Alin Vonica MD, PhD), 1:1000; monoclonal mouse-APOBEC2 (clone 15E11, homemade), 1:5000; TroponinT clone JLT-12 (T6277, Sigma-Aldrich), 1:500; alpha-tubulin DM1A (Abcam, ab7291), 1:5000; MyHC MF-20 (DSHB), 1:20; rabbit anti-SP1 (Merck, 07-645), 1:1000; and rabbit anti-HDAC1 antibody (ab7028), 1:2000.

For co-immunoprecipitation experiments, 4 x 10^6^ C2C12 cells were plated and lysed after 4 days in differentiation medium. Cells were trypsinized, washed with cold PBS, and lysed in 1 mL cell lysis buffer: 0.5% Tween 20, 50 mM Tris pH 7.5, 2 mM EDTA, and freshly added 1X Halt protease and phosphatase inhibitor cocktail EDTA-free (Thermo, 78441). Mixture was vortexed and incubated on ice for 10 min, twice. Nuclei were separated from the cytoplasmic fraction by centrifugation (6000 g, 1 minute, 4°C). Nuclei were washed with 1 mL cold PBS before lysing in 250 μL high salt nuclear lysis buffer: 800 mM NaCl, 1% NP40 (Igepal CA-640), 50 mM Tris pH 7.5, and freshly added 1X protease and phosphatase inhibitor cocktail, EDTA-free. Mixture was vortexed and incubated on ice for 10 min, twice. Nuclear lysate was then diluted to a final salt and detergent concentration of 400 mM NaCl and 0.5% NP40 using 250 μL dilution buffer: 50 mM Tris pH 7.5 and 1X protease and phosphatase inhibitor cocktail, EDTA-free. Nuclear lysates were treated with Benzonase (Merck-Millipore, 70664). Nuclear lysates were pre-cleared on 25 μL Dynabeads M-280 Sheep anti-mouse or antirabbit IgG (Thermo, 11201D/12203D), depending on primary IgG antibody. Pre-cleared nuclear lysate was then added to 50 μL beads conjugated with 2-4 μg primary IgG antibody: rabbit anti-APOBEC2 (Sigma, HPA017957) or rabbit IgG isotype control. Immunoprecipitation was done overnight at 4C with rotation. Beads were thoroughly washed before resuspending and boiling in SDS-PAGE sample buffer.

### Immunofluorescence staining and fusion index of C2C12 cells

C2C12 cells (5×10^4^ cells) were seeded in collagen-coated coverslips (BD Biosciences, 356450) in 12-well plate the day before inducting of differentiation with 2% horse serum. They were washed with cold PBS and fixed with paraformaldehyde (4%) in PBS for 10 minutes at 4 °C. This was followed by 2 washes, 5 minutes each at room temperature and blocking solution (0.5% BSA, 1% gelatin, 5% normal goat serum, 0.1% Triton) in PBS for 1 hour at room temperature. This was followed by overnight stain with antibodies in a humidified chamber at 4°C, three washes with cold PBS 5 min each at room temperature. Coverslips were then incubated with secondary antibodies for 1 hour at room temperature, followed by three washes with PBS 5 min at room temperature. Immunofluorescence staining of C2C12 cells was carried out with primary antibodies: MyHC MF20 (DSHB) and FLAG M2 (Sigma, F1804). Nuclei were counterstained and coverslips were mounted with VECTASHIELD Antifade Mounting Medium with DAPI (Vector Laboratories, H-1200). Images were taken using confocal Leica TCS SP5 II or widefield Zeiss Cell Observer and image analysis was done with Fiji/ImageJ.

### Chromatin salt-extraction profiling

C2C12 cells were seeded in equal numbers (2×10^6^ cells) and induced to differentiate after 12 hours. Five days after differentiation they were lysed in the plate with 100 μl sucrose lysis buffer (320 mM sucrose, 0.5% NP-40, 10 mM Tris pH 8.0, 3 mM CaCl_2_, 2 mM Mg acetate, 0.1 mM EDTA). Extracts were incubated for 5 minutes on ice and spun at 500 g for 5 minutes to collect the nuclear pellet and supernatant as the cytosol fraction. Nuclear pellets were washed with no-salt Nuclei Buffer (50 mM Tris pH 8, 1% NP-40, 10% glycerol). Following the washes the nuclear proteins were extracted at increasing concentrations of NaCl from 250 mM up to 2 M in Nuclei Buffer during which they are homogenized using dounce tissue grinders (Fisher, K8853000000), incubated on ice for 10 minutes and spun at 4°C for 10 additional minutes. Eluted material was collected, resolved on polyacrylamide gel electrophoresis (Criterion XT Bis-Tris Gel 12%, Bio-Rad) and immunoblotted with specific antibodies: Histone H4 (Merck, 05-858R), 1:5000; monoclonal mouse-APOBEC2 (clone 15E11, homemade), 1:5000; α-tubulin DM1A (Abcam, ab7291), 1:5000.

### RNA expression analysis

Library preparation and sequencing were done by Rockefeller University Genomics Resource Center [https://www.rockefeller.edu/genomics/] using TruSeq Stranded mRNA Sample Prep kit as per manufacturer’s instruction. The procedure includes purification of poly-adenylated RNAs. Libraries were sequenced with 50bp paired-read sequencing on the HiSeq2500 (Illumina). Paired end read alignments and gene expression analysis were performed with the Bioinformatics Resource Center at Rockefeller University. Paired-end reads were aligned to mm10 genome using the subjunc function in the Bioconductor Rsubread^44^ package and bigWig files for visualization were generated from aligned reads using the Bioconductor rtracklayer^45^ and GenomicAlignments packages^46^. For analysis of differential expression, transcript quantifications were performed using Salmon^47^ in quasimapping mode. Gene expression values were calculated from transcript quantifications using tximport^48^. Gene expression changes were identified at a cut off of 5% FDR (benjamini-hockberg correction) using the Wald test implemented in DESeq2^49^. Annotation files used: BSgenome.Mmusculus.UCSC.mm10(v1.4.0);org.Mm.db(v3.5.0); TxDb.Mmusculus.UCSC.mm10.knownGene.gtf.gz(v3.4.0)

### RNA editing analysis

RNA editing analysis was performed as previously reported elsewhere^22^. Editing detection was performed by comparing C2C12 control samples (GFPsh) to APOBEC2 knockdown samples using RNA-Seq datasets in triplicates for each sample. Minimum filters include quality control thresholds (minimum of five reads covering the putative site with at least two reads supporting the editing event; filtering of reads that contain indels or support an edit in the first or last two base pairs of a read). Stringent filters applied to the APOBEC1 dependent C-to-U edited sites include all of the above and additionally the magnitude of the control vector was at least 15 and the angle between the wild-type and knockout vectors was at least 0.11 radians, as described in the paper referenced in this section.

### Enhanced reduced representation bisulfite sequencing (ERRBS)

ERRBS library preparation, sequencing and read alignment was performed by the Epigenomics Core Facility of Weill Cornell Medicine [epicore.med.cornell.edu/] as previously described^50, 51^. The procedure includes bisulfite conversion of the DNA. Libraries were sequenced with 50bp single reads (SR) in HiSeq2500 (Illumina). Reads were aligned to a bisulfite converted reference mouse genome with Bismark^52^. The methylation context for each cytosine was determined with scripts from the core facility.

Here coverage of specific genomic regions by ERRBS dataset, refers to the percent of features (eg percent of promoters, CpG islands) that contain at least one CpG that is well covered (> 10x). For gene specific annotations the mm10 UCSC knownGene annotations from the UCSC table browser were used and for CpG islands the mm10 cpgIslandExt track of the UCSC table browser. Genomic features were defined as: CpG islands, CpG island shores were defined as 2kb upstream and downstream of a CpG island; Gene promoters (region 2kb upstream and 2kb downstream of the TSS), exons, introns and intergenic regions.

### Differential methylation analysis

MethylKit (v1.3.8)^53^ was used to identify differentially methylated cytosines (DMCs) with q-value less than 0.01 and methylation percentage difference of at least 25% after filtering ERRBS dataset by coverage, normalizing by median and including CpG sites that are covered >10x, in 3 out of 5 biological replicates (lo.count = 10, lo.perc = NULL, hi.count = 1000, hi.perc = 99.9), (destrand=TRUE,min.per.group=3L). eDMRs (v0.6.4.1)^54^ was used to empirically determine differentially methylated regions, using the DMCs identified with methylKit. In order for a region to be defined as a DMR, default parameters (num.DMCs=1, num.CpGs=3, DMR.methdiff=20) of eDMR were used, so that each region has: (1) at least 1 DMC in the region, as determined using methylKit, (2) at least 3 CpGs included in the region and (3) absolute mean methylation difference greater than 20%. For a region to be defined as a significant DMR, default parameters were used (DMR.qvalue = 0.001, mean.meth.diff = 20, num.CpGs = 5, num.DMCs = 3) so that each significant DMRs has (1) 5 CpGs where at least 3 of them are significant DMCs as determined by methylKit (2) have a minimum 20% methylation change for the region.

### Chromatin immunoprecipitation method

C2C12s were plated at ~70% confluence 12 hours prior to inducing differentiation (seed ~2×10^6 cells) maintained in DMEM (ATCC, 30-2002) with 10%FBS. This was followed by media change to DMEM with 2% horse serum (Life Biotechnologies, 26050-088) to induce differentiation. The cells (~5×10^6 /10cm plate) were harvested at 24-hour or 34-hour after inducing differentiation. They were fixed on plate with 1% PFA in PBS for 10 minutes at RT. Glycine was added to a final concentration of 125mM. Cells were washed 2 times with 1x PBS with protease inhibitor cocktail (Roche, 11836170001). They were lysed on the plate with cold Farnham lysis buffer to ~10×10^6 cells /mL (5mM PIPES pH 8.0, 0.5% NP-40, 85mM KCl,1mM EDTA, PIC) and incubated rotating for 15min at 4°C. Lysates were scraped off the plates, pelleted and resuspended in LB2 (10 mM Tris pH 8.0, 200 mM NaCl, 1 mM EDTA, 0.5 mM EGTA, PIC) and incubated rotating for 15 minutes at 4°C and then centrifuged. Pellets were resuspended to 5×10^7 cells/mL in LB3 (10 mM Tris pH 8.0, 100 mM NaCl, 1 mM EDTA, 0.5 mM EGTA, 0.1% sodium-deoxycholate, 0.5% sodium lauroyl sarcosinate, PIC) until suspension was homogenized. Samples were then sonicated using Covaris ultrasonicator model S220 for 15 minutes with the following settings: 140W peak power, 5% duty, 200 cycles per burst. TritonX-100 to a final concentration of 1% was added to the samples. Samples were clarified by centrifugation at 20,000 g for 10 minutes at 4°C. The supernatant is the soluble chromatin extract. The soluble fragmented chromatin from ~2.5×10^7 was used for each IP. For each IP 100ul Dynabeads (Thermofisher anti-rabbit M280, 11203D) were mixed with 10ul polyclonal rabbit-APOBEC2 antibodies (gift from Alin Vonica MD, PhD) incubating overnight (~16 hours). A magnetic stand was used to separate beads from the lysate and beads were washed one time each with for 5min in: low salt wash (0.1%SDS, 1%Triton X-100, 2 mM EDTA, 20 mM Tris pH8, 150 mM NaCl, PIC), high salt wash (0.1%SDS, 1% Triton X-100, 2mM EDTA, 20mM Tris pH8, 500mM NaCl, PIC), lithium chloride wash (150mM LiCl, 1% NP-40, 1% NaDOC, 1mM EDTA, 10mM TrispH8, PIC), TE wash (10mM Tris-HCl pH8, 1mM EDTA, 50mM NaCl, PIC). Beads were resuspended in 52 ul of elution buffer (50mM Tris-HCl pH8, 10mM EDTA, 1%SDS) and incubated at 30min at 65°C while shaking to prevent beads from settling. The eluate was transferred to a new tube, inputs of the same volume were incubated for 8 hours at 65°C to reverse the crosslink. The samples were treated with RNAse (Roche, 11119915001) for 1 hour at 37°C, and with Proteinase K for 2 hours at 55°C. Fragmented DNA was purified with Ampure beads (Agencourt AMPure XP beads A63881) as per the manufacturer’s instructions.

### Chromatin immunoprecipitation sequencing and analysis

The ChIP-Seq included biological triplicates for each group. ChIP-Seq libraries were prepared using NEBNext Ultra DNA Library Prep Kit as per manufacturer’s instructions. Libraries were sequenced with 75 base pair single read sequencing on the NextSeq 500 (Illumina). Read alignments and initial analysis were performed with the Bioinformatics Resource Center at Rockefeller University. Single-end reads were aligned to mm10 genome using the subread function in the Bioconductor Rsubread^44^ package and bigWig files for visualization were generated from aligned reads using the Bioconductor rtracklayer^45^ and GenomicAlignments packages^46^. Quality metrics for the ChIP-Seq data were assessed using ChIPQC bioconductor package^55^, according to Encyclopedia of DNA Elements (ENCODE) working standards and guidelines for ChIP experiments^56^. Reads mapping to more than one genomic location were filtered prior to peak calling using Model-based Analysis of ChIP-Seq (MACS2)^57, 58^ with duplicate filtering applied and input DNA sample as a control. Peaks that are reproducible (present in 2 out of 3) were filtered for known artifact or blacklisted regions (The ENCODE Project Consortium, 2012). For each of the peaks a weighted mean location of peak summits cross biological replicates is calculated^59^. The list of binding regions 100 base pairs around the mean peak summits was used for downstream analysis. Ngs.plot (v2.61) was used with specific parameters (-G mm10 -D refseq -C -L 1000 -FL 150 -P 4 -SC 0,1 -GO none -RB 0.05) to generate average profiles of ChIP-Seq reads (Shen et al., 2014). ChIPSeeker (v1.14.2)^60^ and ChIPpeakAnno (3.12.7)^61, 62^ were used for downstream analysis after peak calling for annotation of the binding regions to the nearest gene. We created an APOBEC2 occupied gene set, using genes that show consistent APOBEC2 occupancy at both 14-hour and 34-hour time points. GSEA (v3.0)^63^ was used for testing the enrichment of the APOBEC2 occupied gene set in the list of genes that are differentially expressed. Annotation files used: BSgenome.Mmusculus.UCSC.mm10 (v1.4.0) org.Mm.db (v3.5.0) and TxDb.Mmusculus.UCSC.mm10.knownGene.gtf.gz(v3.4.0).

### Gene list analysis

Gene list analyses either by statistical overrepresentation test or statistical enrichment test were done through PANTHER^64^. Briefly, gene lists were filtered based on expression (log2FoldChange, up- or downregulated at specific treatment and time point) and p-adj values (FDR< 10%) and used as input in PANTHER gene list analysis. For statistical overrepresentation tests of upregulated genes with A2 vs GFP shRNA, genes were filtered based on log2FoldChange > 0.58 and FDR < 10% at each time point and used as input list with Mus musculus (all genes in database) as reference/background list. Default parameters were followed for the analyses and are indicated in the corresponding output. For

### Prediction of binding motifs

Transcription factor motifs associated to 108 TF modules^29^ were mapped against time-point specific sequences harboring APOBEC2, 200 base pairs centered on peak summits. For each time point, we defined a background set of negative sequences using scrambled regions of the positive sequences. Using both sequence sets of positives and negatives, we assessed the presence of strong 8-mers associated to each of those 108 families and their ability to classify between APOBEC2 regions and negative sequences, summarizing a Receiver-Operating Characteristic Area Under the Curve (ROC-AUC) for each of those. Assessment of significance in each case was done using a Wilcoxon rank sums test (one sided). *P*-values were corrected through a Benjamini-Hochberg procedure.

### Enrichment of ChIP-seq peaks on APOBEC2 differentially expressed target genes

Using the ChIP-Atlas as a reference, we downloaded all datasets associated to myoblast or C2C12 cells (*N*=54). For each dataset, we intersected ChIP-seq peaks from APOBEC2 in each timepoint and replicate (command *fisher* in *bedtools)*, obtaining a 2 x 2 contingency matrix. The number of overlaps was linked to the closest gene using 2000bp with respect to TSS annotations in the mouse genome. The proportion of genes associated to a differential expression comparison was done by dividing the number of APOBEC2 peaks proximal to a DE-gene with a peak from a ChIP-Atlas dataset by the total number of DE-genes in that comparison. This was repeated for each gene expression contrast. Mean log2 fold change estimates for each ChIP-Atlas peak dataset were obtained by calculating the distribution of log2 fold changes between non-target DE-genes and target DE-genes, in each time point, using the three APOBEC2 ChIP-seq replicates. With those, we calculated a global mean and standard deviation across all ChIP-Atlas factors, reporting a Z-score for dataset, time point and differentially expression comparison between APOBEC2 target and non-target genes.

### BioID samples preparation

Constructs encoding mouse APOBEC2 fused to BirA-Flag by the N- and C-terminus were cloned into pMX-puro. Both constructs, as well as the eGFP-BirA-Flag control, were modified to encode a weak Kozak sequence (TATTGT<u>ATG</u>) to reduce protein expression levels. Virions containing the pMX vectors were produced in the Plat-E packaging cell line^65^. C2C12 cells were spininfected with Plat-E supernatant, 8mg/ml polybrene (16,000 rpm, 90 min, 30C) and selected with 4 μg/ml puromycin to obtain stable cell populations. Similar bait and control expression levels were confirmed by western blot, and localization of the APOBEC2 constructs analyzed by immunofluorescence. C2C12 cells expressing each construct were pre-cultured for 24 h with 2% low horse serum before supplementing the media to 50 μM biotin (BioBasic). Cells were harvested 24 h later, washed 3× with PBS, then lysed in 1.5 mL of RIPA buffer and sonicated 30 s at 30% amplitude (3 × 10 s bursts with a 2s break in between). Benzonase (250 U, Sigma) was added to the lysates during centrifugation, 30min at 16,000×*g*, 4°C. Forty μL aliquots of supernatant were kept to monitor expression and biotinylation and run on western blot, and the remaining lysate was incubated with 70 μL of pre-washed streptavidin-sepharose beads (Sigma) for 3 h on a rotator at 4 °C. Beads were then washed with 1 mL of RIPA buffer, transferred to a new tube and washed again 2× with 1 mL of RIPA buffer and then 3× with 1 mL of 50 mM Ammonium Bicarbonate (ABC) (Biobasic). Beads were then resuspended in 100 μL of ABC with 1 μg of trypsin (Sigma) and incubated overnight at 37 °C with rotation. The following day, 1 μg of trypsin was added for a further 2 h digestion. Samples were centrifuged 1 min at 2000 RPM, and the supernatant was transferred to a new tube. Beads were rinsed twice with 100 μL of water, and all supernatants were pooled and adjusted to 5% formic acid. Samples were then centrifuged for 10min at 16,000×*g* for clarification. Trypsin-digested peptides in the supernatant were dried in a SpeedVac (Eppendorf) for 3h at 30 °C. Samples were resuspended in 15 μL of 5% formic acid and kept at −80 °C for mass spectrometric analysis.

### BioID MS Data analysis

Mass spectrometry was performed at the IRCM proteomics platform. Samples were injected into Q Exactive Quadrupole Orbitrap (Thermo Fisher), and raw files were analyzed with the search engines Mascot and XTandem!^66^ through the iProphet pipeline^67^ integrated in Prohits^68^, using the mouse RefSeq database (version 73) supplemented with ‘‘common contaminants’’ from the Max Planck Institute (http://maxquant.org/downloads.htm), the Global Proteome Machine (GPM; http://www.thegpm.org/crap/index.html) and decoy sequences. The search parameters were set with trypsin specificity (two missed cleavage sites allowed), variable modifications involved Oxidation (M) and Deamidation (NQ). The mass tolerances for precursor and fragment ions were set to 15 ppm and 0.6 Da, respectively, and peptide charges of +2, +3, +4 were considered. Each search result was individually processed by PeptideProphet^69^, and peptides were assembled into proteins using parsimony rules first described in ProteinProphet^70^ using the Trans-Proteomic Pipeline (TPP). TPP settings were the following: -p 0.05 -x20 -PPM -d “DECOY”, iprophet options: pPRIME and PeptideProphet: pP.

### BioID interactions scoring

Six biological replicates of each bait and paired eGFP controls were done in two independent experiments (3 replicates each) and combined for the analysis for maximal statistical power. The estimation of interactions scorings was performed for proteins with iProphet protein probability ≥ 0.9 and unique peptides ≥ 2, by combining two algorithmic approaches: SAINTexpress^71^ and DESeq^72^. The SAINTexpress (version 3.6.1) analysis was performed with default settings using no compression for controls or baits. Interactions displaying a BFDR ≤ 0.01 were considered statistically significant. We also used DESeq2 (version 1.2.1335), an R package that applies negative binomial distribution to calculate enrichments over controls. DESeq was run using default settings and significant preys were selected by applying a ≤0.1 p-value cut-off. The combined list of significant preys obtained from DEseq and SAINT was defined as potential APOBEC2 proximity interactors.

### BioID annotations and network analyses

Graphical representations of protein networks were generated with Cytoscape^73^ (version 3.8.2). Prior to the importation of APOBEC2’s network in Cytoscape, mouse to human orthologs were extracted from the Ensembl database with the BioMart export tool (Mouse genes version GRCm38.p6). Next, we extracted human prey-prey interactions from BioGrid release 4.2.192^74^ and from Cytoscape’s PSICQUIC built-in web service client (April 2021 release) by searching against the IntAct database^75^. Once augmented, clusters were extracted with the Markov CLustering Algorithm (MCL) from Cytoscape’s ClusterMaker2 application (version 1.3.1)^76^. To identify relevant complexes among clusters, the APOBEC2 interactome was annotated with the Gene Ontology Annotation database^77^ (GOA version 171) and CORUM (version 3.0)^78^, a database of known protein complexes. EnrichmentMaps^79^ of GO Biological Processes were generated by importing g:Profiler’s^80^ generic enrichment file outputs and mouse GO BP GMT file. p-values ≤ 0.05 and q-value ≤ 0.05 were set as node cutoffs, and Edge cutoff (similarity) were set at 0.345.

### DNA editing detection

We aligned all short reads from input and IP experiments to the mouse genome (GRCm38 EnsEMBL 90) using HiSAT v2.1.0 with default settings. We removed all non-unique mappers and marked all read duplicates with picard.sam.markduplicates.MarkDuplicates (v 2.5.0). We compared all samples to the reference genome using JACUSA v1.2.4 in call-1 mode with program parameters: call-1 -s -c 5 -P UNSTRANDED -p 10 -W 1000000 -F 1024 -- filterNM_1 5 -T 1 -a D,M,Y -R. Diverging positions are reported if the LLR ratio exceeds 1.0. Briefly, read count distributions at every genomic position (coverage >5) are contrasted with the expected read count based on the reference base. For the pairwise comparison of all input samples stratified by conditions, we used JACUSA v.1.2.4 in call-2 mode with program parameters: call-2 -s -c 5 -P UNSTRANDED,UNSTRANDED -p 10 -W 1000000 -F 1024 -- filterNM_1 5 --filterNM_2 5 -T 1 -a D,M,Y -u DirMult:showAlpha -R. Briefly, read count comparison from replicate input samples are contrasted with one another: A2 shRNA knockdown vs GFP shRNA knockdown. Diverging positions are reported if the LLR ratio exceeds 1.0.

### Recombinant mouse APOBEC2 production

Recombinant His_6_-tagged mouse APOBEC2 proteins were produced in Sf21 insect cells by the EMBL Protein Expression and Purification Core Facility. The genes encoding mouse APOBEC2 were cloned into the pFastBac HTa vector (Thermo) and these constructs were used for transposition into E. coli DHIOEMBacY cells (Geneva Biotech). The isolated bacmid DNA was utilised for generation of the recombinant baculoviruses. For the mouse APOBEC2 protein production, 5 mL of baculovirus was used to infect 1 L of Sf21 cells at a density of 1 x 106 cells/ml. After 72h, the cells were harvested by centrifugation (30 min, 600 x g, 4°C) and resuspended in lysis buffer (20 mM Tris pH 8.0, 800 mM NaCl, 20 mM imidazole and 5 mM β-mercaptoethanol) supplemented with benzonase, 10 mM MgCl_2_ and cOmplete EDTA-free protease inhibitors (Roche). The cells were lysed using a Dounce homogenizer and the lysate was cleared by centrifugation (30 min, 20000 x g, 4°C). The cleared lysate was loaded onto a 5 mL Ni-NTA column (Macherey-Nagel). After washing with a buffer consisting of 20 mM Tris pH 8.0, 300 mM NaCl, 20 mM imidazole and 5 mM β-mercaptoethanol, the Ni-NTA column was eluted using a gradient up to 300 mM imidazole. The elution fractions containing mouse APOBEC2 were pooled and dialysed overnight at 4°C to ion exchange buffer (20 mM Tris pH 8.0, 100 mM NaCl, 1 mM DTT). The dialysed sample was loaded onto a 5 mL HiTrap Heparin HP column (Cytiva) coupled to a 5 mL HiTrap Q HP anion exchange column (Cytiva). After washing, the HiTrap Heparin HP column and the HiTrap Q HP column were eluted separately in a gradient ranging from 100 mM to 1M NaCl. Finally, the mouse APOBEC2 protein eluted from the HiTrap Q HP column was subjected to a size exclusion chromatography (SEC) step using a HiLoad 16/600 Superdex 75 pg column (Cytiva) preequilibrated with SEC buffer (20 mM HEPES pH 7.5, 150 mM NaCl and 0.5 mM TCEP). When removal of the His_6_-tag was required, His_6_-tagged TEV protease (produced in-house) was added to the purified mouse APOBEC2 protein. After the overnight TEV cleavage step at 4°C, Ni-NTA beads (Qiagen) were added to the sample and incubated for 1h at 4°C. After centrifugation (1 min, 100 x g, 4°C), untagged mouse APOBEC2 was recovered from the flow through of the Ni-NTA beads. Recombinant mouse APOBEC2 proteins were aliquoted, flash-frozen with liquid N2 and stored at −80°C.*Microscale Thermophoresis (MST)*

Purified APOBEC2 was labeled using Cy5 Mono NHS Ester (GEPA15101, Sigma-Aldrich) at 5 mg/mL protein and a 3:1 dye:protein ratio. Labeling reactions were performed in 20 mM HEPES, pH 7.5, 150 mM NaCl, and 0.5 mM Tris(2-carboxyethyl)phosphine (TCEP). Reactions were incubated overnight at 4°C with constant agitation. After incubation, reactions were deactivated using quencher buffer (ab102884, Abcam). Remaining dye was washed away by concentrating protein using Vivaspin^®^ 500 centrifugal concentrators (Sartorius) into MST buffer (10 mM HEPES pH 7.5, 50 mM NaCl, 5 mM MgCl_2_). The degree of labeling was typically at about 1.

Oligonucleotides (oligos) were ordered from Sigma-Aldrich: SP/KLF motif F: GGC GGC GCG GCC CCG CCC CCT CCT CCG GC; SP/KLF motif R: GCC GGA GGA GGG GGC GGG GCC GCG CCG CC; A-tract motif F: TCT CAA GAA AAA AAA AAA AAG AC; A-tract motif R: GTC TTT TTT TTT TTT TCT TGA GA.

Oligonucleotides were annealed by incubating at 95°C and slow cooling to 25°C before storing at 4°C. Oligonucleotides were diluted to MST buffer + 0.05% Tween-20 for the final reaction. MST buffer was supplemented with 0.05% Tween-20. Cy5-labeled APOBEC2 was held constant at 50 nM while the oligonucleotides were titrated (1:1 dilution) between 19.53 to 20,000 nM. Reactions were incubated for 30 min before loading into standard glass capillaries (MO-KO22, NanoTemper Technologies). MST measurements were performed using a Monolith NT.115 (NanoTemper Technologies) at 85% LED power and 40% MST power. Data represent 3 independent measurements. MO.Affinity Analysis v2.1.3454 (NanoTemper Technonolgies) was used for curve fitting and calculating Kd values with Thermophoresis + T Jump settings.

### Electromobility shift assay (EMSA)

Oligos used for MST were labeled with [γ-^32^P] ATP and annealed with complementary oligos to form double-stranded (ds) oligos. Specified recombinant protein, mouse APOBEC2 and/or human SP1 (Sigma, SRP2030), and ss or ds oligos were mixed in binding reaction buffer (10 mM HEPES pH 7.5, 50 mM NaCl, 5 mM MgCl_2_, 5% glycerol and 5 mM DTT) for 30 minutes at room temperature. Reactions were resolved on 5% TBE gel (3450048, BioRad) with 0.5X TBE buffer (1610733, Bio-Rad) for 1.5 hours at 120 volts with the tank submerged on ice. Gels were then dried and imaged with a phosphorimager system (Azure Biosystems, Inc.).

## Acknowledgements

The authors thank Dr. Diego Mourao-Sa for his insights and support during the completion of these studies; Dr. Thomas Carroll, for reproducing RNA-Seq and ChIP-Seq bioinformatics analyses; Dr. Pete Stavropoulos, Erik Debler, Philipp Schmiege and Nicholas Economos for initially producing APOBEC2 protein and mouse monoclonal (15e11) antibody; Dr. Frank Schwarz and GPCF@DKFZ for assistance with the MST experiments. We would like to thank Julia Flock and Dr. Kim Remans (Protein Expression and Purification Core Facility, EMBL) for producing the recombinant mouse APOBEC2 in Sf21 insect cells. Last but not least, FNP and LM would like to thank Dr Bruce McEwen (1938-2020) and Dr Karen Bulloch. Without their lasting support and encouragement this work would not have been completed.

## Author Contribution

LM and JPL designed the study, performed experiments, analyzed data, and wrote the manuscript with supervision from FNP. SR for performing supplementary experiments. DH performed RNA editing analysis. ILR performed motif prediction and enrichment of ChIP-seq peaks analysis with supervision from JBZ. PGS and JR performed experimental work and JB performed analysis related to BioID data with supervision from JMDN. AV provided key reagent. CD performed DNA editing detection analysis. All authors wrote, read, and approved the final manuscript.

---

**Supplementary Figure 1.**
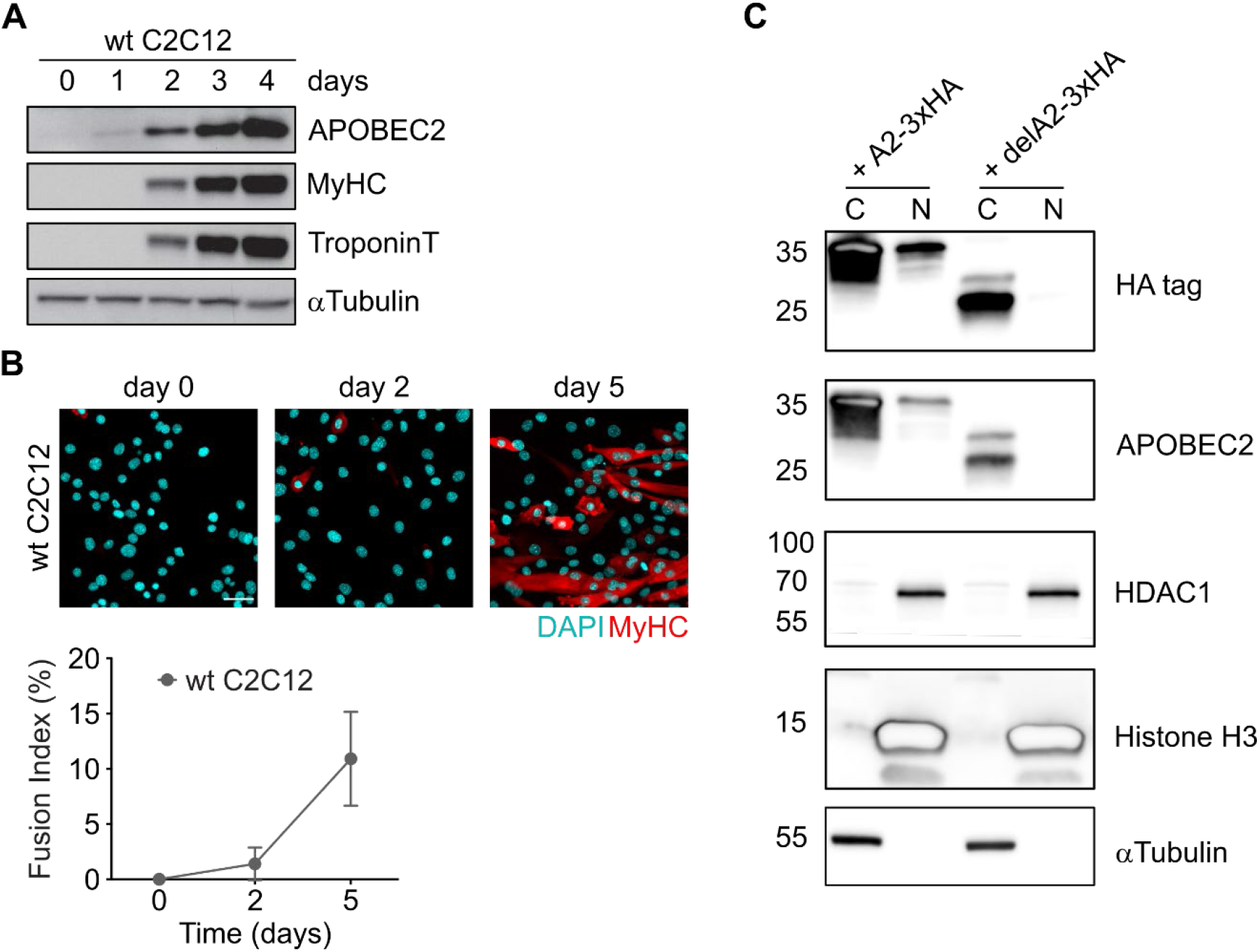
C2C12 myoblast differentiation. **(A)** Whole cell extracts of mouse wildtype (wt) C2C12 myoblasts and myotubes were analyzed by Western blotting using anti-APOBEC2 antibodies. MyHC and TroponinT were used as markers of late differentiation, alpha-tubulin was used as loading control. **(B)** C2C12 cells were cultured in differentiation medium for 0, 2 and 5 days, fixed, and stained with antibody to MyHC (red). Nuclei were visualized by DAPI staining (blue). Below the quantification of differentiation expressed as fusion index, which is the percentage MyHC-positive myotubes with >2 nuclei. Results are presented as means from quantification of at least 6 images/sample. Error bars indicate SD. Scale bar = 50 μm. **(C)** Cytoplasmic and nuclear (C and N respectively) were taken from C2C12 cells transduced with vectors to overexpress 3xHA-tagged wildtype APOBEC2 (+ A2-3xHA) and truncated APOBEC2 (+ delA2-3xHA). Western blots were performed with respective lysates using antibodies against HA-tag, APOBEC2, HDAC1, Histone H3, and αTubulin.

**Supplementary Figure 2.**
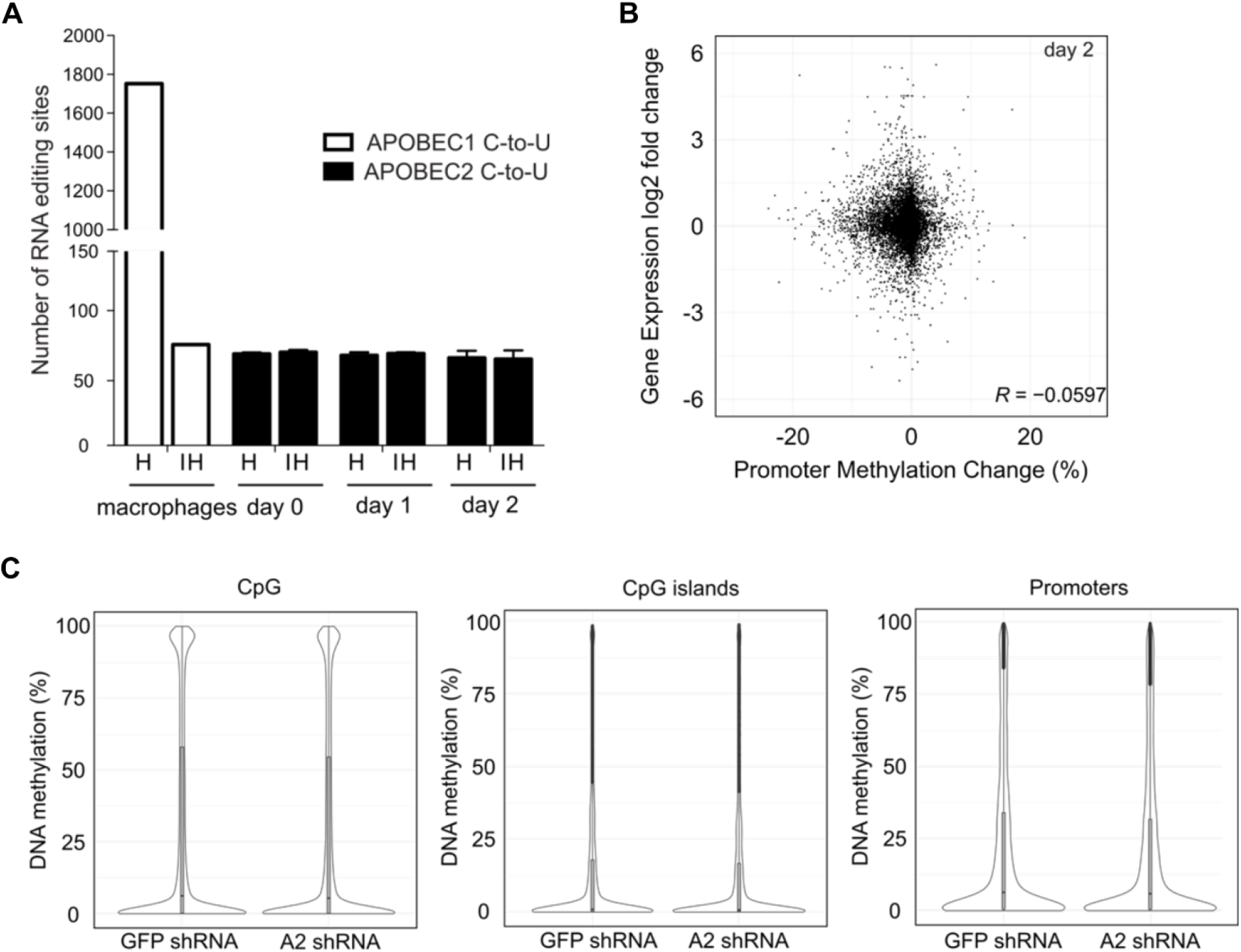
mRNA editing and DNA demethylation in C2C12 myoblasts. **(A)** Candidate C-to-U RNA editing sites called from APOBEC2 knockdown samples, control (GFPsh) at day 0, 1, and 2 in DM in C2C12s and wild-type and APOBEC1-/- macrophages (positive control). Hits (H) represent candidate editing sites present in control (GFPsh) but not in APOBEC2 knockdown dataset (in positive control hits are the # of sites in APOBEC1-/- dataset.) Inverse hits (IH) represent putative edited sites yielded when the inverse comparison is made, thus edit sites present in the APOBEC2 knockdown dataset but not in the control (GFPsh) (for the positive control edit sites that are present in APOBEC1-/- dataset but not in wild-type). Data are represented as means ± SD using outputs of 3 RNA-Seq datasets. **(B)** Methylation changes across all the represented promoters in the ERRBS dataset compared with the expression changes of the same genes in RNA-Seq dataset. Shown here are datasets from the day 2 timepoint. R = Pearson’s correlation coefficient. **(C)** Distribution of DNA methylation frequencies in C2C12s as determined by ERRBS for individual CpGs, CpG islands and promoters. Promoters are defined at -/+ 2Kb around the TSS in Ensemble annotations. CpG islands were taken from the cpgIslandExt track of the UCSC table browser. Violin plots represent the distribution of DNA methylation frequencies for each feature. Median and first and third quartiles are shown with the box plots.

**Supplementary Figure 3.**
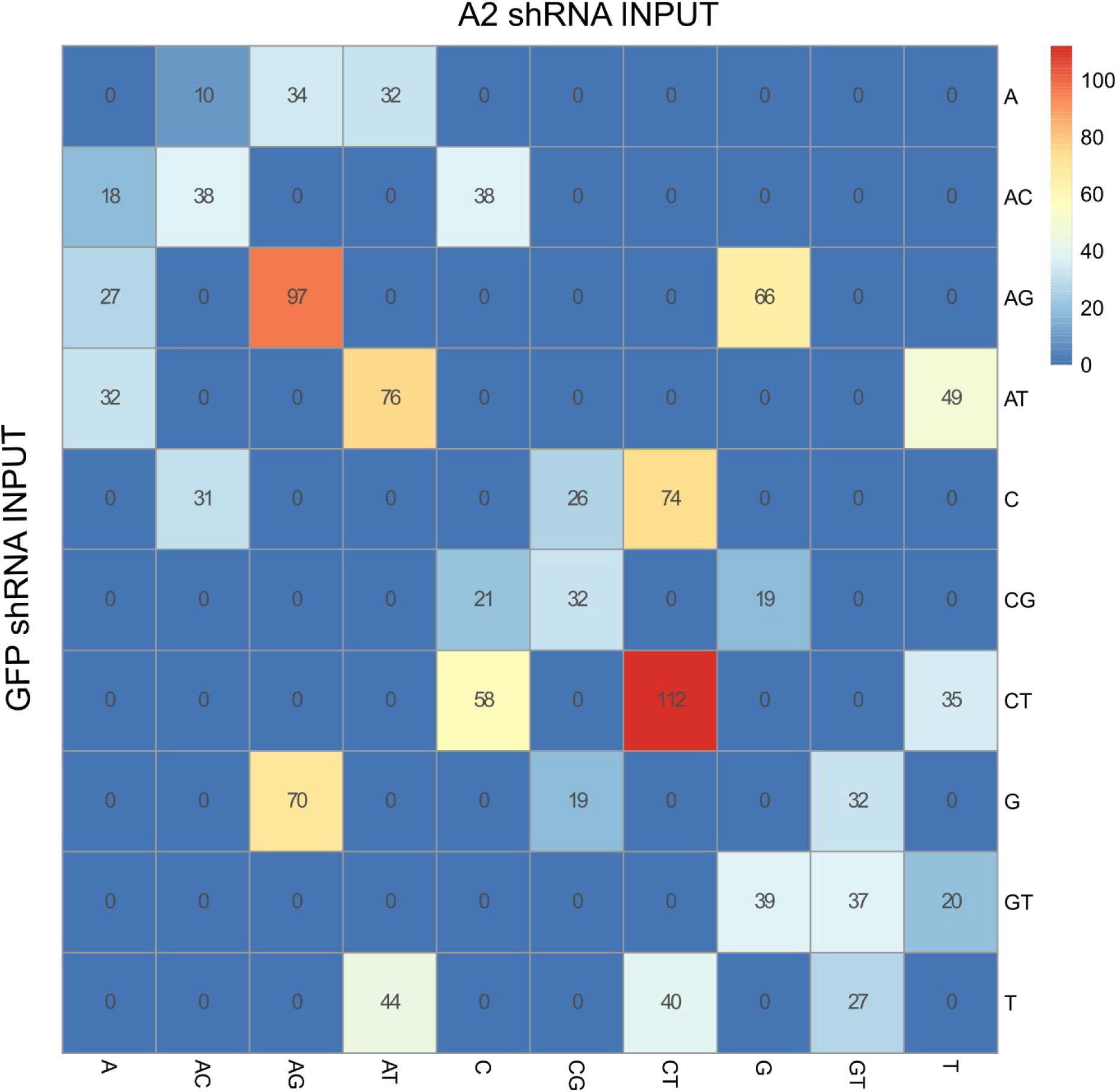
DNA Editing in APOBEC2 knockdown versus control (GFP shRNA) Pairwise heatmap - This is a head-to-head comparison of variant positions between the 2 input sample sets. The symmetry of the heatmap indicates that there is no preference for a unidirectional base substitution process.

**Supplementary Figure 4.**
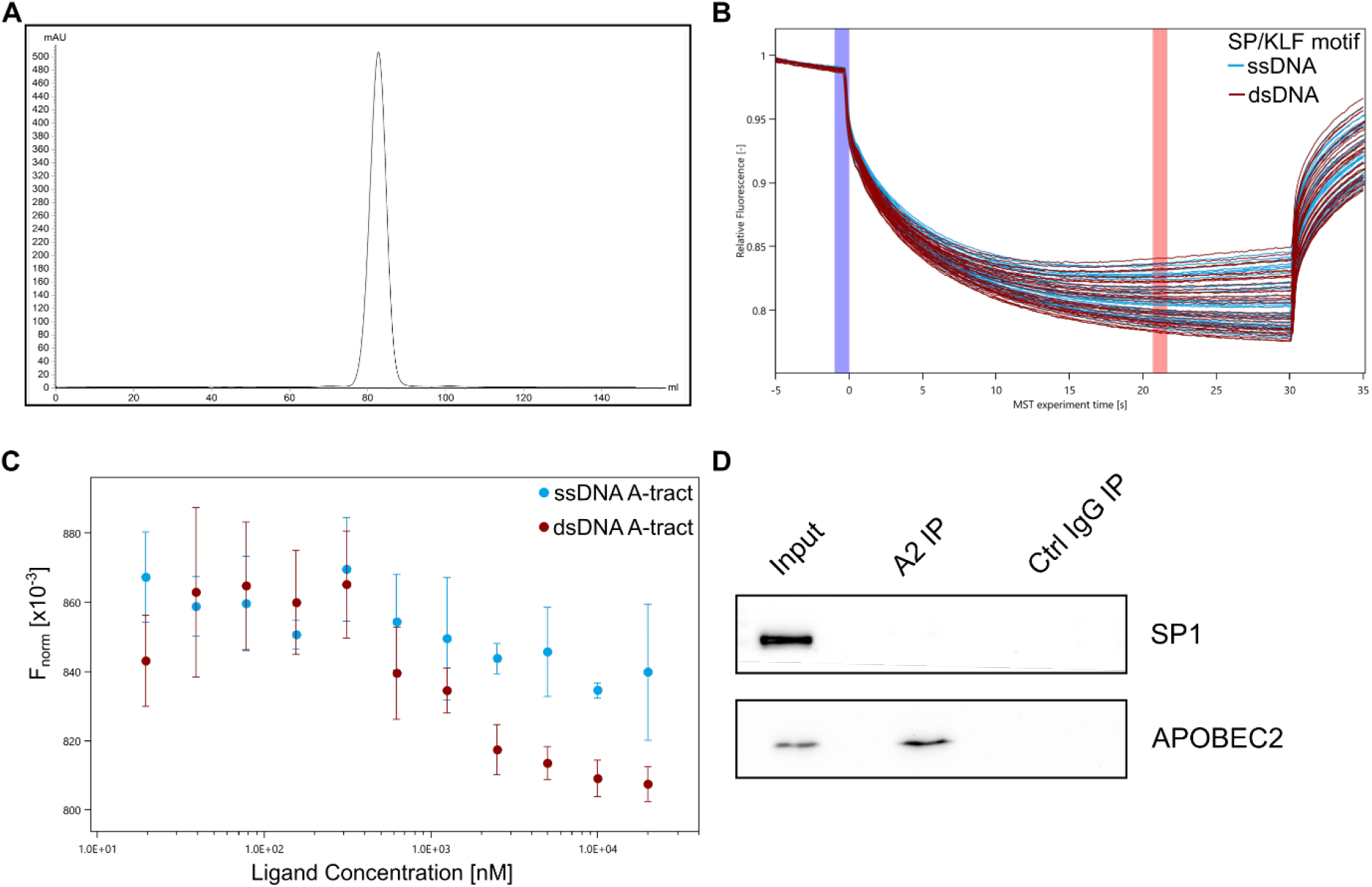
Recombinant APOBEC2 electromobility shift assays. **(A)** Size exclusion chromatogram (Superdex 200, GE Healthcare) of recombinant 6x-His-APOBEC2 produced in High Five/Sf9 (Thermo) insect cells. **(B)** MST trace for APOBEC2 interaction with ssDNA and dsDNA SP/KLF motifs. Traces correspond to experiment in Figure 4A. Blue and red highlighted regions represent cold and hot regions respectively that were used for the standard thermophoresis+ T-jump analysis. **(C)** MST experiments measuring purified APOBEC2 binding to single-stranded DNA (ssDNA) or double-stranded (dsDNA) with A-tract motif. Cy5-labeled APOBEC2 was kept constant (50 nM) while the concentration of non-labeled SP/KLF motif was titrated (1:1 dilution) between 20 nM – 20,000 nM. Normalized fluorescence (Fnorm) values and graph were produced with MO.Affinity Analysis v2.1 (NanoTemper Technonolgies). Values represent 3 independent measurements with error bars representing the standard deviation. **(D)** Co-immunoprecipitation (Co-IP) of APOBEC2 with SP1 in C2C12 myoblasts differentiated to myotubes for 4 days. Nuclear protein lysates (Input) were incubated with beads conjugated to either APOBEC2 antibody (A2 IP) or IgG isotype control antibody (Ctrl IgG IP). Proteins were then eluted, ran on an SDS-PAGE gel, and blotted with APOBEC2, or SP1 antibodies.

**Supplementary Figure 5.**
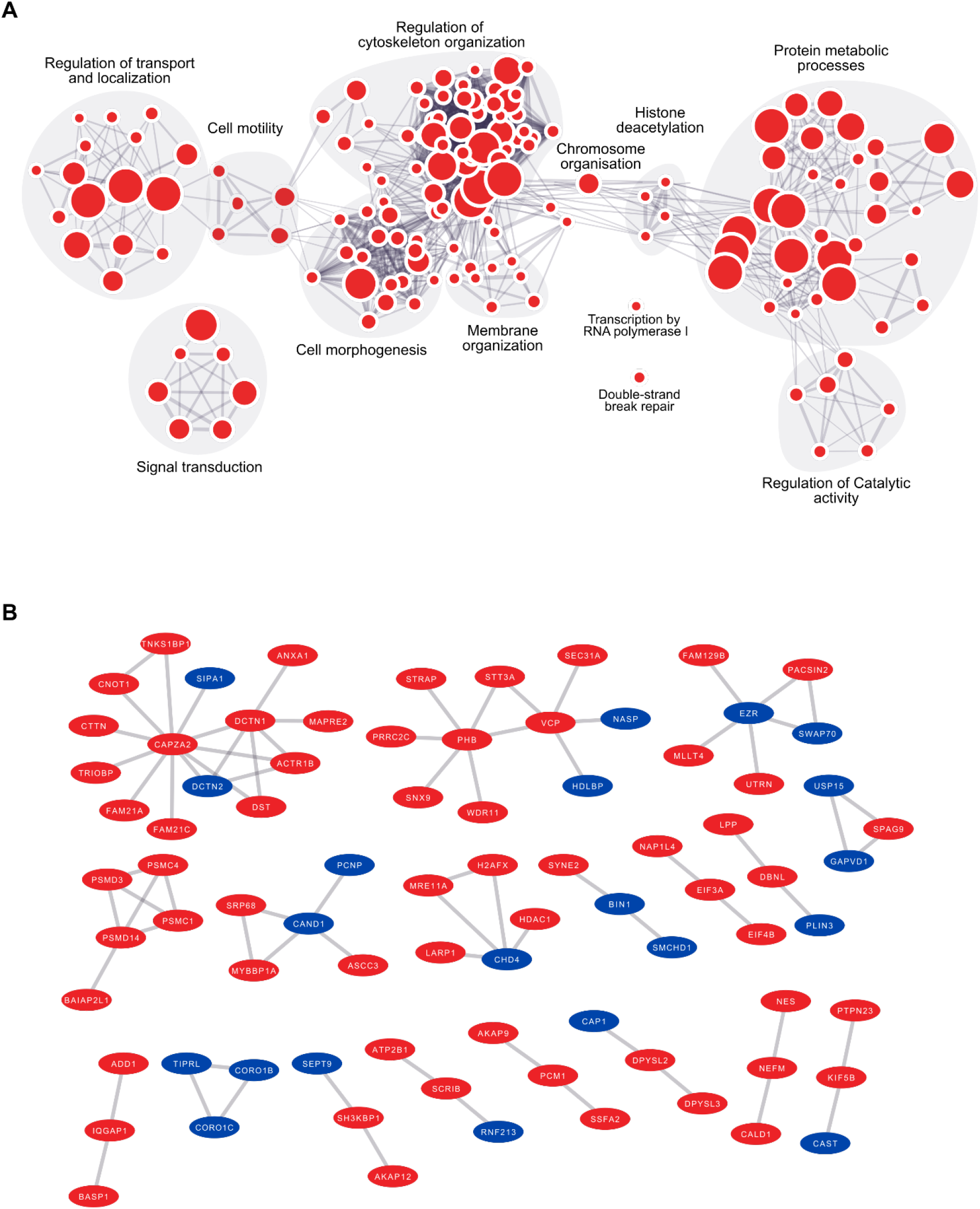
BioID GO Enrichment Map. **(A)** Gene ontology enrichment map based on APOBEC2 BioID hits annotated by gProfiler (BFDR<0.05). Gene ontology terms related to similar biological processes are clustered (indicated in grey). **(B)** Protein complexes identified by proximity-dependent protein biotinylation (BioID) in C2C12 expressing APOBEC2-BirA or BirA-APOBEC2. Each red node corresponds to a protein that was significantly enriched in either or both A2-expressing cells compared to GFP-BirA control cells. Nodes in blue indicate proteins that were preferentially labeled by APOBEC2-BirA compared to AID-BirA in mouse B lymphocytes in a previously published dataset^82^. The edges denote the known interactions of these proteins with each as described in Methods.

